# Bacterial and fungal contributions to delignification and lignocellulose degradation in forest soils with metagenomic and quantitative stable isotope probing

**DOI:** 10.1101/387308

**Authors:** Roland C. Wilhelm, Rahul Singh, Lindsay D. Eltis, William W. Mohn

## Abstract

Delignification, or lignin-modification, facilitates the decomposition of lignocellulose in woody plant biomass. The extant diversity of lignin-degrading bacteria and fungi is underestimated by culture-dependent methods, limiting our understanding of the functional and ecological traits of decomposers populations. Here, we describe the use of stable isotope probing (SIP) coupled with amplicon and shotgun metagenomics to identify and characterize the functional attributes of lignin-, cellulose-and hemicellulose-degrading fungi and bacteria in coniferous forest soils from across North America. We tested the extent to which catabolic genes partitioned among different decomposer taxa; the relative roles of bacteria and fungi, and whether taxa or catabolic genes correlated with variation in lignocellulolytic activity, measured as the total assimilation of ^13^C-label into DNA and phospholipid fatty acids. We found high overall bacterial degradation of our model lignin substrate, particularly by gram-negative bacteria (Comamonadaceae and Caulobacteraceae), while fungi were more prominent in cellulose-degradation. Very few taxa incorporated ^13^C-label from more than one lignocellulosic polymer, suggesting specialization among decomposers. Collectively, members of Caulobacteraceae could degrade all three lignocellulosic polymers, providing new evidence for their importance in lignocellulose degradation. Variation in lignin-degrading activity was better explained by microbial community properties, such as catabolic gene content and community structure, than cellulose-degrading activity. SIP significantly improved shotgun metagenome assembly resulting in the recovery of several high-quality draft metagenome-assembled genomes and over 7,500 contigs containing unique clusters of carbohydrate-active genes. These results improve understanding of which organisms, conditions and corresponding functional genes contribute to lignocellulose decomposition.

## Introduction

The incomplete decomposition of woody biomass in coniferous forests is an important global carbon sink, with approximately one third of a gigaton of carbon accruing on an annual basis (Myneni et al., 2001). Lignocellulose decomposition is influenced by the structure and function of microbial communities (Strickland *et al.*, 2009; Cleveland *et al.*, 2014) and terrestrial carbon cycling models increasingly parameterize these properties (Allison *et al.*, 2012; Wieder *et al.*, 2015). However, efforts are constrained by our rudimentary knowledge of the composition and ecology of decomposer populations, stemming from limitations of culture-dependent methods and the complexity of soil communities. The best characterized decomposers inhabit forest soil litter, where conditions favour rapid lignocellulose degradation (Melilo *et al.*, 1989; Cortrufo *et al.*, 2015), yet the decay of lignin-rich plant polymers in soil occurs in a continuum governed by conditions and substrate accessibility, resulting in diversified niches for decomposers (von Lützow *et al.*, 2006; Lehmann and Kleber, 2015; Klotzbücher *et al.*, 2015). To resolve the catabolic and ecological traits of decomposers, we must utilize culture-independent methods, like stable isotope probing (SIP), that better reflect *in situ* conditions.

The nature of microbial lignin-degradation is poorly described beyond the canonical breakdown of lignin in woody biomass by specialized, aerobic, litter-inhabiting wood-rot fungi.Yet, these fungi are largely absent in deeper mineral soil where conditions favour decomposition by bacteria (Ekschmitt *et al.*, 2008), who represent the most likely lignin-degraders when oxygen is limited (Benner *et al.*, 1984; DeAngelis^a^ *et al.*, 2011; Hall *et al.*, 2015). Several soil bacteria can degrade model lignin compounds in pure culture, suggesting a role for them in the catabolism of low-molecular weight, partially-degraded forms of lignin (Crawford, 1978; Masai *et al.*, 2007; DeAngelis^b^ *et al.*, 2011; Brown *et al.*, 2012; Taylor *et al.*, 2012; Davis *et al.*, 2013; Rashid *et al.*, 2015). Knowledge of bacterial lignin-degradation in environmental contexts is limited, with all studies of soil degraders originating from tropical forests, which describe active populations of predominantly Alpha-and Gammaproteobacteria (DeAngelis^a^ *et al.*, 2011; Pold *et al.*, 2015). The existing evidence for bacterial lignin-degradation demonstrates the need to characterize both bacterial and fungal populations and contrast their roles in different soil environments to better understand the *in situ* processes that govern the decomposition of lignocellulose.

Delignification rapidly increases the rate of lignocellulose decomposition and may have evolved primarily as a means of accessing more readily degradable plant carbohydrates, exemplified by the strategies of white-and brown-rot Agaricomycota (Eastwood *et al.*, 2011; Ding *et al.*, 2012; Zeng *et al.*, 2014). The capability to co-degrade lignin and other lignocellulosic polymers has not been thoroughly explored in bacteria. There is some evidence for the co-metabolism of lignocellulosic polymers by various *Streptomyces spp.* (Větrovský *et al.*, 2014), while a comparative genomics study revealed that cellulose-degrading bacteria also possess higher numbers of hemicellulases (Medie *et al.*, 2012). The extent to which catabolic traits are conserved within specific taxa versus communities will influence models of microbial decomposition. In the simplest case, the abundance of highly adapted, multi-substrate degrading taxa may predict rates of decomposition, which is supported by a small number of studies (Strickland *et al.*, 2009; Wilhelm^a^ *et al.*, 2017). However, forces of genomic streamlining in bacteria lead to functional diversification among closely related species, particularly in extra-cellular processes that produce common goods (Morris *et al.*, 2012), evident in the species-level conservation of bacterial endoglucanases (Berlemont and Martiny, 2013). A multi-substrate SIP experiment provides the means to identify whether forest soil decomposers can assimilate ^13^C from various lignocellulosic polymers and, with shotgun metagenomic sequencing, determine whether their genomes encode the necessary suite of catabolic genes.

To address the above knowledge gaps, we utilized SIP microcosm-based experiments to investigate the composition and degradative potential of hemicellulose-, cellulose-and lignin-degrading populations from organic and mineral layer soils in coniferous forests across North America. The identity of degraders and their genomic content were assessed using amplicon and shotgun sequencing of DNA enriched in ^13^C from labeled substrates.Lignocellulolytic activity was quantified according to the amount of ^13^C assimilated into total DNA and phospholipid fatty acids (PLFA). The *in situ* abundances of lignocellulolytic populations were determined on the basis of previously reported pyrotag libraries from the same field samples (Wilhelm *et al.*, 2017 ^b^, Wilhelm *et al.*, 2017^c^). This is currently the most comprehensive cultivation-independent study of lignocellulolytic populations in forest soils and yields new insights to the taxa responsible and the importance of certain catabolic gene families.

## Material and Methods

### Overview of Sites and Sample Collection

Soil samples were collected from five forest regions in North American chosen to encompass a range of climates, tree cover and soil conditions. Samples were collected from mature coniferous forest corresponding to ‘reference’ plots in the Long-Term Soil Productivity Study in northern Ontario (BS_ON_ and JP_ON_), British Columbia (IDF_BC_), California (PP_CA_) and Texas (LP_TX_). Detailed descriptions of geographical locations, soil properties and sampling methods can be found in Wilhelm^b^ *et al.* (2017) as well as in Table S1. Samples were collected from three sites in each geographical region, except IDF_BC_ where three plots were samples from a single site. The organic layer (O-horizon) and the top 20 cm of mineral soil (A-horizon + occasionally upper B-horizon) were sampled separately (overview in Figure 1a). Samples were kept at 4°C during transport, sieved through 2-mm mesh and stored at −80°C.

**Figure 1.**
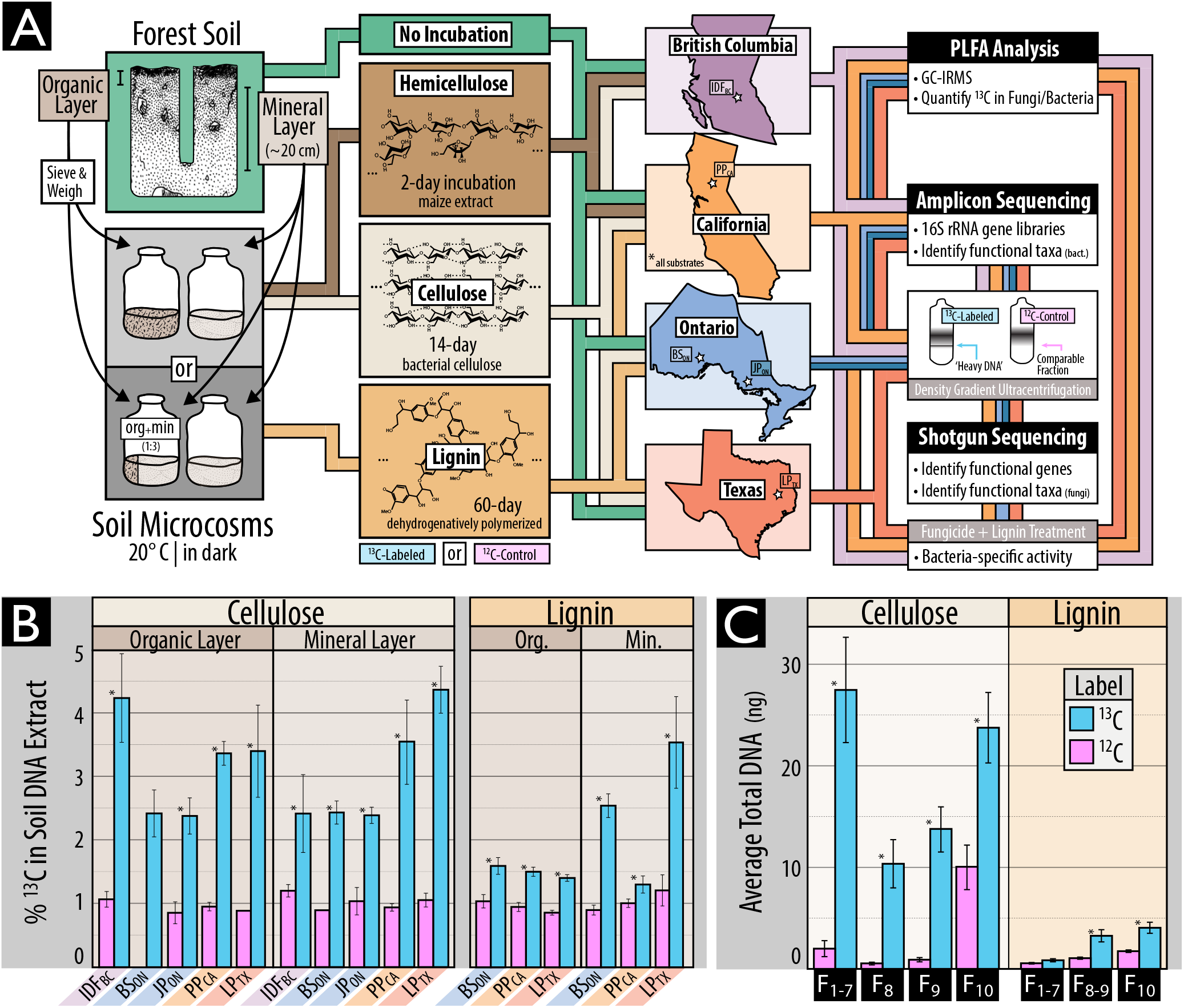
(A) An overview of the samples and methods used in this study, including (B) evidence of the enrichment of ^13^C in DNA extracted from soil amended with ^13^C-labeled or unlabeled substrate, and (C) for the separation and recovery of ^13^C-labeled DNA from heavy fractions of a cesium chloride density gradient. The schematic in A illustrates our use of organic and mineral layer soil from five forest regions incubated with or without one of three model constituents of lignocellulose. Due to limited quantities of ^13^C-labeled substrate, the complete pairing of all forest regions and substrates was not possible. In B and C, Statistically supported differences between paired labeled and unlabeled treatments (t-test; *p* < 0.01) are designated with an asterisk (*).

### Preparation of SIP Soil Microcosms and ^13^C-labeled Substrates

Microcosms were prepared with a minimum of three replicates by adding ^13^C-labeled hemicellulose, cellulose or lignin to soil from as many of the five regions as possible contingent on the availability of substrate (Figure 1a). Microcosms were comprised of either 1 g (organic) or 2 g (mineral) dry wt soil in 30-mL serum vials adjusted to a moisture content of 60% (mineral) and 125% (organic) (w/v). Microcosms were wetted and pre-incubated in the dark at 20° C for one-week prior to the addition of 1% (w/w) ^13^C-labeled cellulose or 0.8% (w/w) lignin. The hemicellulose-amended incubations were performed as part of a previous study examining the impacts of timber harvesting on decomposition using identical methods as described here for cellulose (Leung *et al.*, 2016). Each microcosm with ^13^C-labeled substrate was paired with an identical ‘^12^C-control’ microcosm amended with the corresponding unlabeled substrate (~1.1% atom ^13^C). ^12^C-control microcosms were used to estimate the natural abundance of ^13^C in PLFAs and to control for the background presence of GC-rich DNA in higher density CsCl gradient fractions (Youngblut *et al.*, 2014).

Bacterial cellulose was produced from *Gluconacetobacter xylinus* str. KCCM 10100 grown with either unlabeled or ^13^C-labeled glucose (99 atom *%* ^13^C, Cambridge Isotope Laboratories, MA, USA) in Yamanaka medium as described in the Supplementary Methods. DHP lignin was synthesized from ring-labeled coniferyl alcohol using horseradish peroxidase as described Kirk and Brunow (1988). Coniferyl alcohol was synthesized from ring-labeled vanillin (75 atom % ^13^C, Sigma Aldrich, CA) according the reactions in Figure S1, following methods described in the Supplementary Methods. DHP lignin had a weight average molecular weight (M_w_) of 2,624 g.mol^−1^ which equates to ~ 14 polymerized units of coniferyl alcohol. Yields were approximately 12% and 18% (w/w) and were labeled at ~99% and 60% atom % ^13^C for cellulose and lignin, respectively.

Microcosms were incubated at 20° C for 14 days (cellulose) or 60 days (lignin) or 2 days (hemicellulose; Leung *et al.*, 2016). The incubation length for this study was optimized by preliminary time-course experiments described in Wilhelm *et al.* (2014). To overcome poor ^13^C-labeling of microbial biomass in organic soils from DHP-lignin, organic soils were mixed 1:3 with corresponding double-autoclaved mineral soil to dilute pre-existing organic matter. Following incubation, soil samples were lyophilized and stored at −80° C until processing. All SIP-PLFA, SIP-pyrotag and SIP-metagenomic data were generated from the same microcosm unless otherwise stated. Due to limited quantities of ^13^C-substrate, SIP-lignin microcosms were prepared with soil from only BS_ON_, PP_CA_ and LP_TX_, while SIP-hemicellulose data was available for IDF_BC_ and PP_CA_ (see Table S2 for summary of data). To determine the lignolytic activity solely by bacteria, eight additional soil microcosms were prepared with ^13^C-lignin amended mineral soil and incubated with or without weekly additions of two fungicides: 1 mg.g^−1^ soil cylcoheximide plus 1.2 mg.g^−1^ soil fungizone (amphotericin B).

### SIP-Phospholipid Fatty Acid Analysis

PLFAs were extracted from 0.75 g (organic) or 1.0 g (mineral) dry wt soil according to Bligh and Dyer (1959) and ^13^C-content was analyzed using ion ratio mass spectrometry (UBC Stable Isotope Facility) ported with gas chromatography as detailed in Churchland *et al.* (2013). Peak identification was based on retention time compared against two reference standards: bacterial acid methyl-ester standard (47080-0; Sigma–Aldrich, St. Louis) and a 37-Component fatty acid methyl-ester mix (47885-U; Sigma–Aldrich, St. Louis). For details on the treatment of unidentifiable, but ^13^C-labeled, peaks and taxonomic designations consult the Supplementary Methods.

### Preparation of Amplicon and Shotgun Metagenome Libraries

DNA was extracted from soil (0.5 g) using the FastDNA™ Spin Kit for Soil (MPBio, Santa Ana, CA) following the manufacturer’s protocol. ^13^C-enriched DNA was separated using cesium chloride density centrifugation per methods outlined by Neufeld *et al.* (2007) with minor modifications (see Supplementary Methods). DNA from heavy fractions (1.727-1.735 g. mL^−1^; typically F1-F7) was pooled, desalted and concentrated using Amicon Ultra-0.5 mL filters (EMD Millipore, MA, USA). DNA from ^12^C-controls was treated identically; however, the total DNA recovered from those heavy fractions of ^12^C-controls was insufficient for sequencing library preparation, resulting in the need to pool additional fractions (F1-F9). The % atom ^13^C of soil DNA extracts and DNA from heavy fractions was quantified according to Wilhelm *et al.* (2014). All other DNA quantitation was performed using Pico-Green fluorescent dye (ThermoFisher, MA, USA).

PCR amplification was performed on bacterial 16S rRNA gene (V1–V3) using barcoded primers as described in Hartmann *et al.*, (2012). PCR reactions were performed in triplicate and pooled prior to purification. Samples were sequenced using the Roche 454 Titanium platform (GS FLX+) at the McGill University and Genome Québec Innovation Centre, yielding an average of 8,400 bacterial reads per sample with a minimum length of 250-bp following quality processing. All 16S rRNA gene libraries are stored at the European Nucleotide Archive (ENA) under the study accession PRJEB12502. Identically prepared 16S rRNA gene amplicon libraries for SIP-hemicellulose (Leung *et al.* 2016) and for corresponding unincubated field soil samples (Wilhelm^c^ *et al.* 2017) were re-analyzed in the present study, following retrieval from the ENA.

Whole shotgun metagenomes were prepared from ^13^C-enriched or ^12^C-control DNA using either the Nextera DNA Sample Preparation Kit (Illumina Inc., CA, USA) for SIP-cellulose microcosm (40-50 ng template DNA), or the Nextera XT DNA Sample Preparation Kit (Illumina Inc., CA, USA) for SIP-lignin (1 ng), due to lower recovery of ^13^C-enriched DNA. The two sequencing library preparations produced very similar metagenomic profiles in a direct comparison (Wilhelm, 2016). Four libraries were multiplexed per lane of Illumina HiSeq 2500 (2 x 100-bp) and were grouped based on similar average fragment sizes based on fragment profiles from a 2100 Bioanalyzer (Agilent Technologies) All raw and assembled shotgun metagenomic data is available under ENA accession PRJEB12502, with an overview of accessioned data available in Table S3.

### Bioinformatic and Statistical Analyses

16S rRNA gene amplicon libraries for all three SIP-substrates were quality filtered and simultaneously processed using mothur according to the Schloss “454 SOP” (accessed November 2015; Schloss *et al.* 2009) and clustered into operational taxonomic units (OTUs) at 0.01% dissimilarity. Taxonomic classification was performed using the RDP Classifier (Wang *et al.*, 2007) with the GreenGenes database for 16S rRNA genes (database gg_13_8_99; August 2013). All OTU counts were normalized to total counts per thousand reads. All related count tables, classifications and sample data is provided as phyloseq objects in the Supplementary Data. Phylogenetic trees were prepared by placing sequences into the pre-aligned SILVA tree (Pruesse *et al.*, 2007; ‘SSURef_NR99_123_SILVA_12_07_15’) using maximum parsimony in ARB (Westram *et al.*, 2011).

Shotgun metagenome libraries from SIP-cellulose and SIP-lignin were quality preprocessed with Trimmomatic (Bolger *et al.*, 2014; v. 0.32) and FastX Toolkit (Gordon and Hannon, 2010; v. 0.7). Individual metagenomes were assembled using Ray-meta (Boisvert *et al.*, 2012). Metagenomes were also composited by location and soil layer and assembled with the low-memory assembler MEGAHIT (Li *et al.*, 2015; v. 1.0.2). Contigs from composited metagenomes were binned into metagenome-assembled genomes (MAGs) using MetaBAT (Kang *et al.*, 2014; v. 0.18.6) based on read mapping with Bowtie2 (Langmead and Salzberg,2012). The quality of draft genome bins was assessed using CheckM (Parks *et al.*, 2015) and the lowest common ancestor (LCA) classification of raw, assembled and MAG data was performed with MEGAN (Huson *et al.*, 2007; v. 5.10.1) based on DIAMOND blastx (Buchfink *et al.*, 2015; v. 0.7.9) searches against a local version of the ‘nr’ NCBI database (downloaded October 2014). MAGs were classified to a specific taxon if > 25% of contigs were uniformly classified at the genus level. ORF prediction was performed by Prodigal (Hyatt *et al.*, 2010; v. 2.6.2) and CAZy encoding ORFs were identified based on BLAST searches against a local CAZy database (downloaded August 19^th^, 2015) and hmmscan searches (Finn *et al.*, 2011; HMMER v.3.1b1) using custom hmms for oxidative, lignolytic gene families (see Supplementary Methods) and hmms provided by dbCAN (Yin *et al.*, 2012). Contigs containing clusters of three or more genes encoding CAZymes (‘CAZy gene clusters’) were recovered using a custom approach described in the Supplementary Methods. CAZyme families attributed xylanase, endoglucanase and ligninase activity can be found in the Supplementary Methods.

Statistics were performed using R v. 3.1.0 (R Core Team, 2014) with general dependency on the following packages: reshape2, ggplot2, plyr (Wickham 2007; Wickham 2009; Wickham 2011), Hmisc (Harrell and Dupont 2015) and phyloseq (McMurdie and Holmes 2014). Where necessary, P-values were adjusted according to Benjamini & Hochberg (1995). The ‘vegan’ R-package (Oksanen *et al.*, 2015) provided tools to calculate, non-parametric multidimensional scaling (‘metaMDS’) on Bray-Curtis dissimilarities (‘vegdist’). A combination of DESeq (Anders and Huber 2010) and indicator species analysis (De Cáceres and Legendre 2009) were used to compare control and labeled microcosms to identify OTUs and taxonomic groups that incorporated ^13^C from the labeled substrate. An OTU or taxon had to be selected by at least one of these methods and be, on average, 3-fold more abundant in ^13^C libraries in at least one geographic region. Random forest classification, implemented in boruta Kursa and Rudnicki, 2010), was used to identify taxa or CAZymes predictive of total ^13^C incorporation into DNA or PLFA. Correlation between features selected by boruta and total ^13^C were subsequently performed using ‘rcorr’ from the R package Hmisc. The relative importance of soil layer, geographic region, total carbon, CAZy composition and community structure in linear models for each substrate was assessed using the R package relaimpo (Grömping, 2006) using the primary and secondary axes of NMDS as measures of community structure and CAZy composition.

## Results

We utilized SIP microcosm-based experiments to investigate the taxonomic identity and catabolic potential of lignocellulolytic populations from coniferous forest soils in five regions across North America. Microcosms contained either organic (top 2 – 5 cm) or mineral layer soil (5 – 20 cm) and were amended with either ^13^C-labeled hemicellulose, cellulose or lignin (Figure 1a). Significant enrichment of ^13^C in soil DNA extracts (Figure 1b) and PLFAs, and the recovery of heavy DNA from higher density gradient fractions (Figure 1c) confirmed the catabolism and assimilation of ^13^C from substrates. The quantification of total ^13^C in PLFA and DNA was in close agreement (r = 0.74, *p* < 0.001). The successful targeting of lignocellulolytic populations was evident in the strong compositional differences in DNA and PLFA profiles from soils amended with ^13^C-labeled substrates compared to controls. A total of 503 OTUs were enriched in ^13^C-amplicon libraries, enabling the determination of which taxa had the capacity to degrade multiple lignocellulosic (Figure 2; full list in Table S4). The enrichment of degradative populations was also apparent in shotgun metagenomes, as evidence by the increased proportion of reads comprising the assembly (Figure S2), the assembly quality (N50_13C_ = 4000 vs. N50_12C_ = 2100; *t* = 2.83, *p* = 0.005), and the recovery of a greater number of MAGs from metagenomes prepared from ^13^C-enriched DNA compared to controls (Figure 3). The high quality of MAGs enabled the characterization of gene content to test our conclusions about individual capacities for catabolizing lignocellulosic polymers.

**Figure 2.**
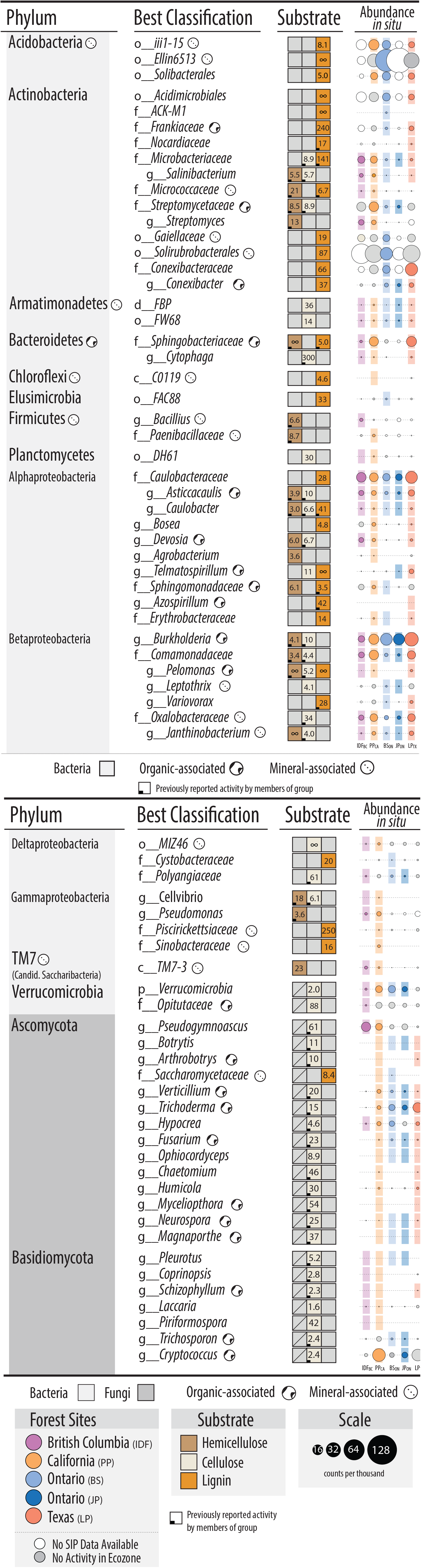
Designations of all putatively hemicellulolytic, cellulolytic and lignolytic taxa based on differential abundance between ^13^C-and ^12^C-pyrotag (bacterial OTUs) or unassembled metagenomes (fungal taxonomy based on LCA). The lowest possible classifications are given with the prefix corresponding to taxonomic rank. The abundance of each taxon *in situ* (i.e. in unincubated field soil) is represented by a scaled circle that was coloured (and given a coloured background) if that taxon was attributed lignocellulolytic activity in that forest region. Uncoloured (white) circles indicate SIP data was not available for that forest region (consult Figure 1a for overview) and grey circles indicate SIP data was available, but the taxon was not attributed activity. The averaged ratio of abundance in ^13^C-versus ^12^C-libraries for each taxonomic classification (i.e. all OTUs within each taxon) is displayed for each substrate. A lemniscate (W) indicates cases where that taxon was detected in only ^13^C-libraries. A full account of all enriched OTUs and citations for all reported catabolic activity is available in Supplementary Table S4.

**Figure 3.**
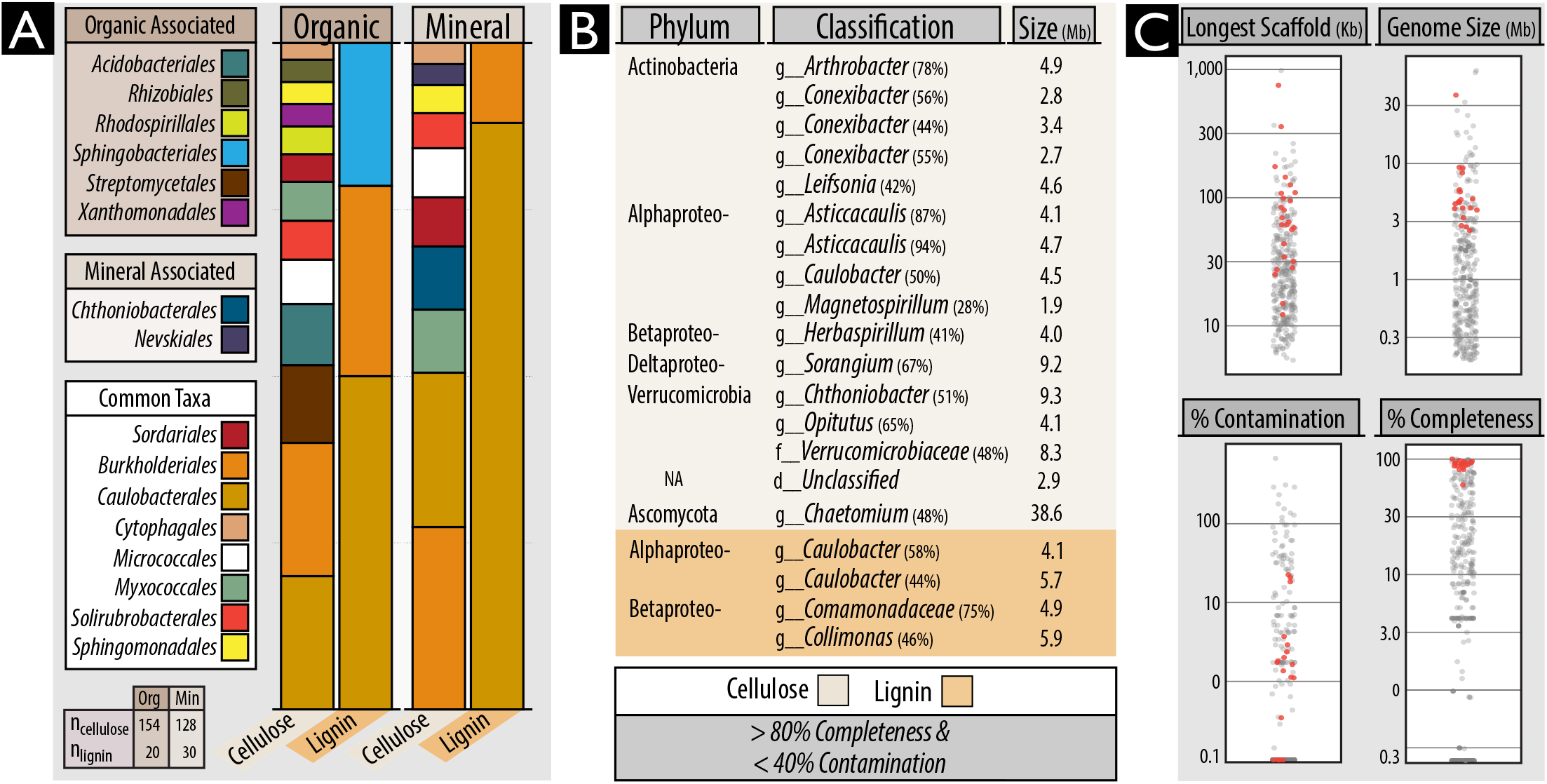
An overview of the classification and quality of metagenome-assembled genomes (MAGs) recovered from organic and mineral soil layers from ^13^C-cellulose and ^13^C-lignin shotgun metagenomes. (A)Proportion of MAGs classified to various Orders. Taxonomic designations are given to any MAG possessing >40% uniformity in the classification of sequence data (normalized to contig length). (B) Information about the top-quality MAGs based on completeness (> 80%) and low contamination (< 40%)based on the completeness or duplication, respectively, in lineage-specific single-copy gene sets. (C) Quality metrics for all MAGs, with top-quality MAGs identified as red points. The complete list and all information on the MAGs is provided in Table S5.

## Identifying Degraders of Multiple Lignocellulosic Polymers

Members of the family Caulobacteraceae and order Burkholderiales dominated ^13^C-enriched DNA pools, comprising between 1 - 5% of ^13^C-metagenomes, and were highly represented in our collection of MAGs (Figure 3). In Burkholderiales, three major families possessed taxa that could degrade one or more substrates: Burkholderiaceae, Comamonadaceae and Oxalobacteraceae (Figure 2; Figure S3). In Caulobacteraceae, members of the genus *Asticcacaulis* assimilated ^13^C from both hemicellulose and cellulose, while members of *Caulobacter* demonstrated the capacity to assimilate all three substrates (Figure 4a; full breakdown in Figure S4). Notably, several OTUs belonging to unclassified clades of Caulobacteraceae were highly enriched in ^13^C-DNA pools from lignin (Figure S5) and were significantly more abundant in mineral than organic soils (21.5 vs. 6.0 counts per thousand reads, respectively; t-test; p=0.02). Differences in predisposition for cellulose-versus lignin-catabolism in *Asticcacaulis* versus *Caulobacter*, respectively, was evident in the catabolic gene content of metagenome assemblies (Figure 4bc).

**Figure 4.**
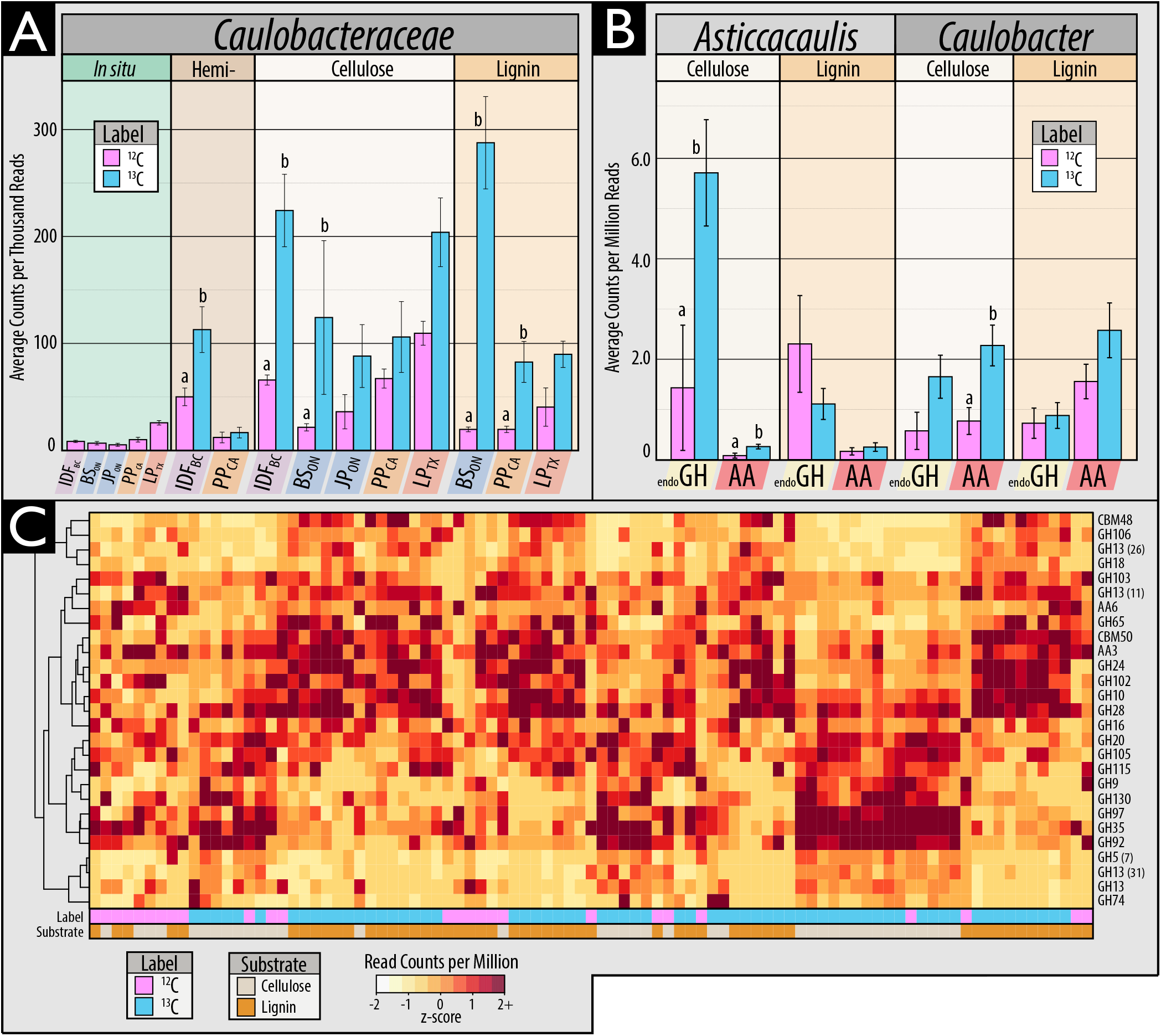
Characteristics of ^13^C assimilation by members of Caulobacteraceae evidenced by (A) theirenriched relative abundance in ^13^C-amplicon libraries for all three substrates; (B) differences in the relative abundance of endoglucanase and AA gene families between closely related genera, *Asticcacaulis* and *Caulobacter*, and (C) a heatmap profiling CAZy gene content in cellulose and lignin metagenome assemblies. In C, the x-axis corresponds to individual metagenomes and the y-axis abundant CAZy families. Both axes are clustered according to Bray-Curtis dissimilarity for the full dataset, though only CAZy families with greater than 5 read counts per million were displayed. Statistically supported differences are grouped by lettering (TukeyHSD; *p* < 0.01). Consult Figure S4 for a complete breakdown of differentially abundant genera in Caulobacteraceae.

We anticipated that lignocellulolytic species would commonly degrade more than one of the three lignocellulosic polymers tested, yet most incorporated ^13^C from only one (Figure 2). No single OTU had increased relative abundance in ^13^C-DNA pools for all three substrates. Only five OTUs were enriched in ^13^C-DNA pools for two substrates and were, in all cases, enriched from ^13^C-lignin and ^13^C-cellulose. These OTUs were classified as *Simplicispira* (Burkholderiaceae), *Aquincola* (Comamonadaceae), unclassified Caulobacteraceae and two Sinobacteraceae (Figure S6). The only genera to possess OTUs that collectively incorporated ^13^C from all three substrates were *Pelomonas* and *Caulobacter.* More taxa possessed OTUs that collectively catabolized a combination of hemicellulose and cellulose (n = 7) than cellulose and lignin (n = 4), or hemicellulose and lignin (n = 3). Genera that utilized both hemicellulose and cellulose included *Cellvibrio*, *Janthinobacterium*, *Cytophaga* and *Salinibacterium*. All fungal taxa enriched in unassembled ^13^C-metagenomes were enriched from cellulose, with the sole exception of a putative member of Saccharomycetaceae from lignin. SIP tended to enrich relatively rare populations, evidenced by the small proportion of OTUs (41/503) shared between ^13^C-amplicon libraries and unincubated field samples representing *in situ* relative abundances.

## Correlating Soil Properties with ^13^C-Assimilating Populations

Variation in the composition of ^13^C-assimilating bacterial populations was explained most by substrate use (PERMANOVA; R^2^ = 13%; *p* , 0.001), followed by the interaction of geographic region and substrate use (10%; *p* < 0.001), region (8%; *p* < 0.001) and the interaction of substrate and site (6%; *p* < 0.001; Table S6). The relative importance of the predictors of ^13^C-incorporation into DNA or PLFA differed for hemicellulose-, cellulose-and lignin-degrading populations (Figure 5a). However, soil layer, forest region and CAZy content consistently ranked highly followed by taxonomic composition and total soil carbon content. Higher soil organic matter content slowed rates of ^13^C-assimilation, with total carbon and nitrogen negatively correlated with ^13^C enrichment of PLFAs in hemicellulose- (ρ = −0.71; *p*_adj_ < 0.01) and lignin-amended mineral layer soils (ρ = −0.85; *p*_adj_ < 0.001; Figure 5c), but not those with cellulose. The highest rates of ^13^C-assimilation were observed for hemicellulose and cellulose in the organic layer of IDF_BC_ and for both cellulose and lignin in LP_TX_ mineral soil (Figure 6a).

**Figure 5.**
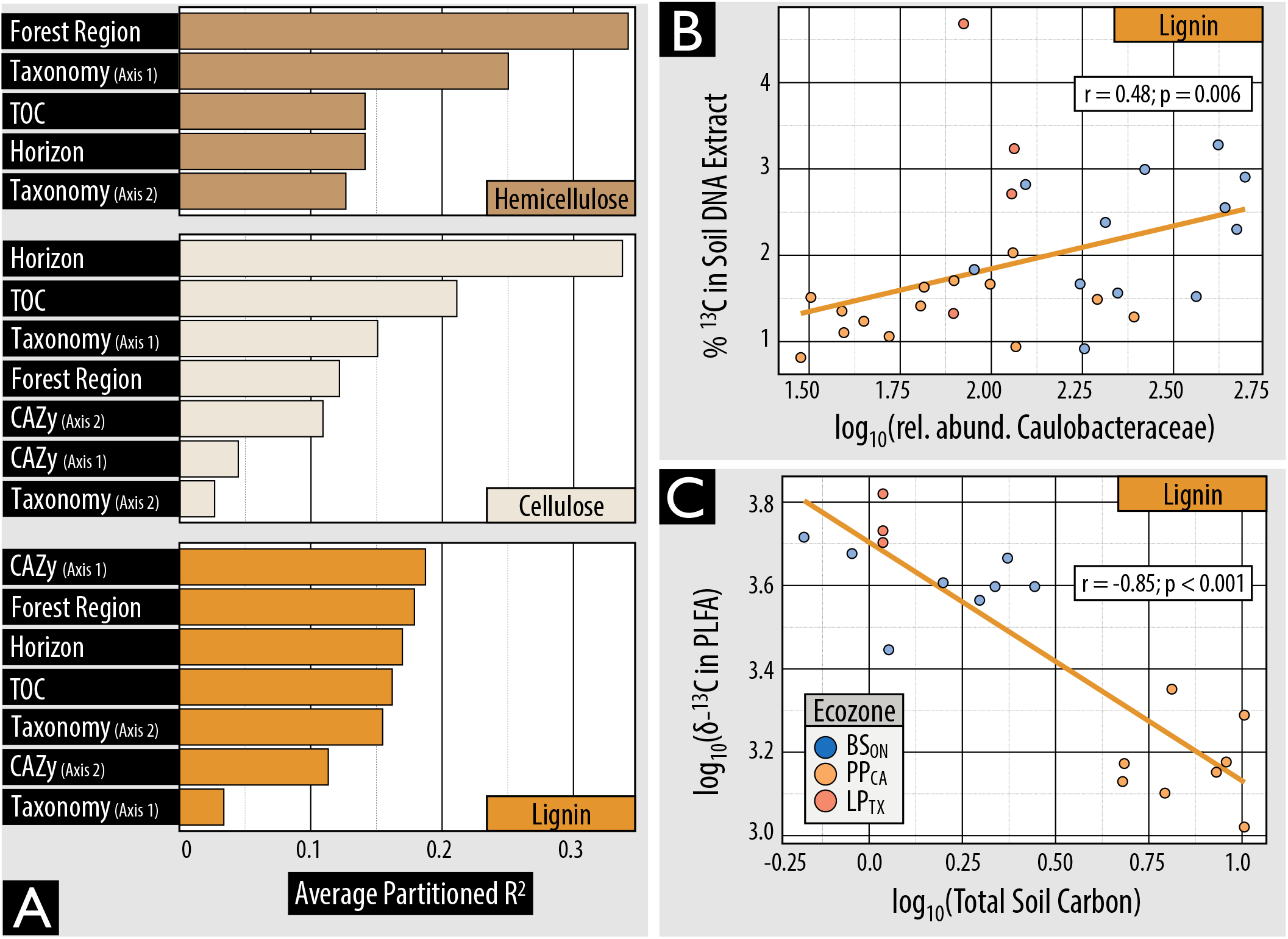
The influence of environmental and microbial community variables on lignolytic activity. (A) Environmental and community compositional variables which explain the highest proportion of variance in total ^13^C incorporation into PLFA. Compositional predictors ‘taxonomy’ and ‘CAZy’ were based on the separation of samples by the primary and secondary axes of NMDS ordinations. (B) The correlation in relative abundance of Caulobacteraceae reads in 16S rRNA libraries and ^13^C-enrichment of DNA for lignin. (C) The negative correlation between total ^13^C incorporation from lignin into PLFA and total soilcarbon in mineral soils. δ- ^13^C is a measure of the ratio of stable isotopes of carbon relative to Vienna Pee Dee Belemnite.

**Figure 6.**
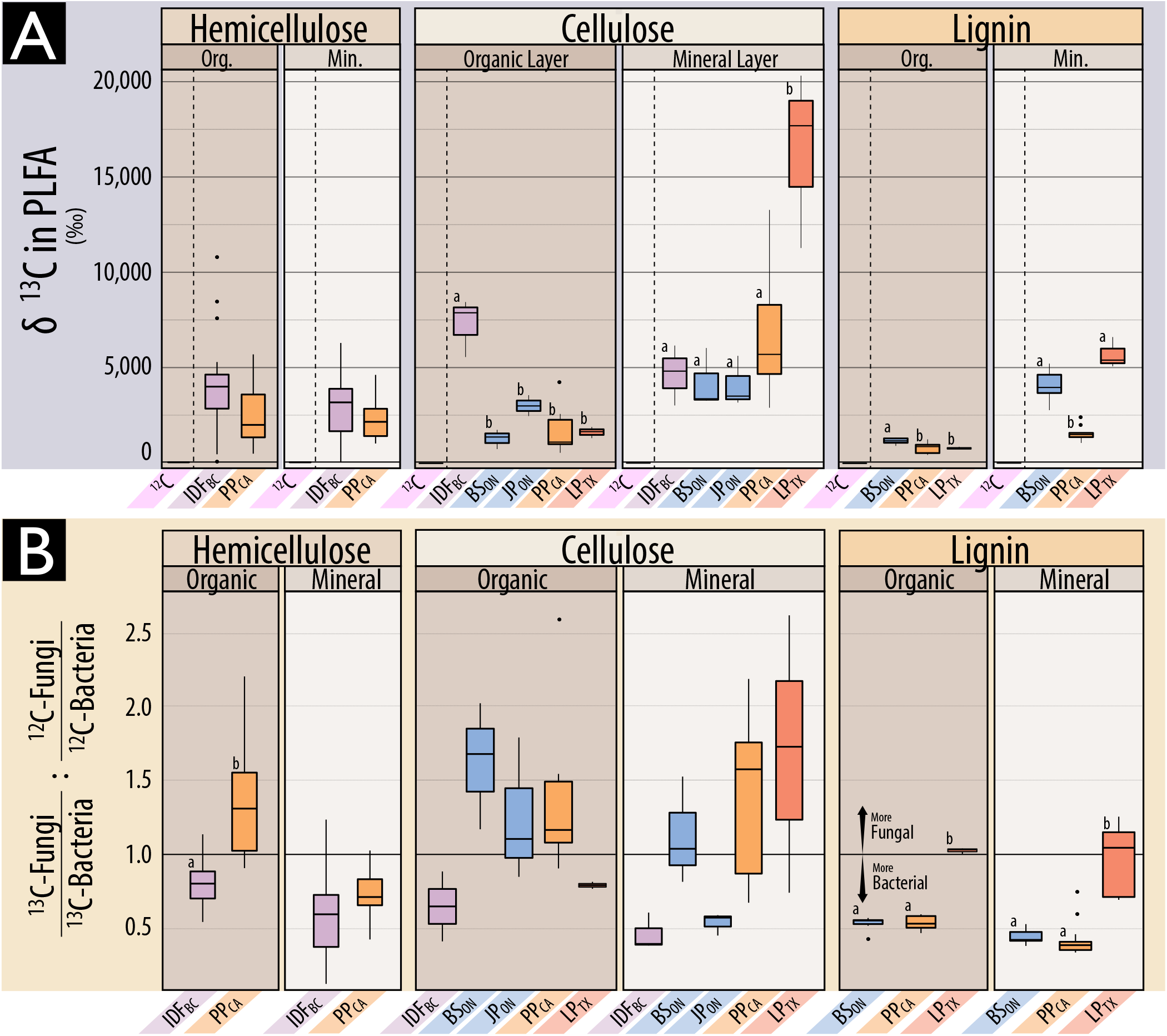
A comparison of ^13^C incorporation into microbial biomass in soils of differing forest region based on (A) δ-^13^C in PLFA, and (B) the relative incorporation into bacterial versus fungal PLFA markers. In A, PLFAs from microcosms incubated with unlabeled substrates had comparable delta-values to soil (−25 ‰), and low variance, and are distinguished by a dashed line. Statistically supported differences are grouped by lettering (TukeyHSD; *p* < 0.01). In B, the ratio represented by the y-axis wasdesigned to normalize ^13^C-enrichment in fungi relative to bacteria (^13^C_fungi_ : ^13^C_bacteria_) to the proportion of pre-existing bacterial and fungal biomass (^12^C_fungi_ : ^12^C_bacteria_). Differences in incubation length of microcosms among substrates should be considered when comparing between them. Also, δ-^13^C enrichment is weighted against total ^12^C, and thus reflects the total lignocellulolytic activity relative to total biomass which is 3- to 20-fold higher in organic layer soils (details in Table S1).

Highly active cellulolytic populations in the IDF_BC_ organic layer were uniquely dominated by bacteria in contrast to those from other regions and soil layers (Figure 6b). These active cellulolytic populations were comprised largely of taxa not active elsewhere, namely: MIZ46 (Deltaproteobacteria), *Cellvibrio* (Gammaproteobacteria), and Planctomyces (vadinHA49) (Figure S7). In contrast, Ascomycota were the major degraders of cellulose in most other soils (i.e. BS_ON_, PP_CA_ and LP_TX_). The active lignolytic populations in BS_ON_ soils were also distinct from other forest soils, and were comprised of Cystobacteraceae (Deltaproteobacteria), FAC88 (Elusimicrobia, formerly Termite Group 1), Gaiellaceae and Solirubrobacterales (Actinobacteria). Twelve bacterial genera were identified by random forest classification whose relative abundance in ^13^C amplicon libraries was significantly positively correlated with ^13^C-enrichment of DNA (Table S7). The strongest correlations occurred for *Caulobacter* (cellulose; ρ = 0.6; p_adj_< 0.001), unclassified Caulobacteraceae (lignin; Figure 5b), and *Janthinobacterium* (cellulose; ρ = 0.59; p_adj_ < 0.001). No fungi were correlated with the ^13^C-enrichment of DNA or PLFA.

## Bacterial versus Fungal Lignocellulolytic Activity

In most microcosms, bacteria were predominantly active in catabolizing and assimilating ^13^C from substrates and potentially other breakdown products (Figure 6b; Figure S8). Gram-negative bacteria had the highest δ-^13^C enrichment of PLFAs in 72% of lignin-amended microcosms, while roughly equal numbers of cellulose-amended microcosms were dominated by gram-negative bacteria or fungi. The total quantity of ^13^C assimilated into PLFAs (μmol.g^−1^ dry wt soil) did not significantly differ between microcosms with greater bacterial versus fungal assimilation (Wilcoxon test; W = 194, p=0.7). Gram-negative and gram-positive bacterial populations dominated the assimilation of hemicellulose in a comparable number of microcosms, 41% and 39%, respectively, though ^13^C-enrichment was 2-fold higher when gram-negative bacteria dominated (Wilcoxon test; W=1261, *p* < 0.001).

Fungicide-treatment greatly reduced assimilation of ^13^C from lignin into fungal PLFAs, but also into gram-positive bacteria and, to a lesser extent, gram-negative bacteria (Figure 7a). While overall bacterial activity was decreased by fungicide-treatment, the assimilation of ^13^C by gram-negative bacteria persisted at relatively high levels (20-50% of untreated incubations) when zero enrichment of fungal PLFAs had occurred (in BS_ON_ and PP_CA_). The major taxa incorporating ^13^C-lignin in fungicide-treated incubations were distinct from those in untreated ones and were predominantly gram-negative bacteria: Burkholderiaceae and Sphingobacteriaceae (Figure 7b). Despite the reduction in ^13^C-enrichment of gram-positive PLFAs, the relative abundance of a major lignolytic Actinobacterial family (Nocardiaceae) and Actinobacteria overall (not shown) were increased in ^13^C-metagenomes after fungicide treatment. The impact of fungicide-treatment on the relative abundance of Caulobacteraceae differed by region, with only minor decreases observed in PP_CA_ and LP_TX_.

**Figure 7.**
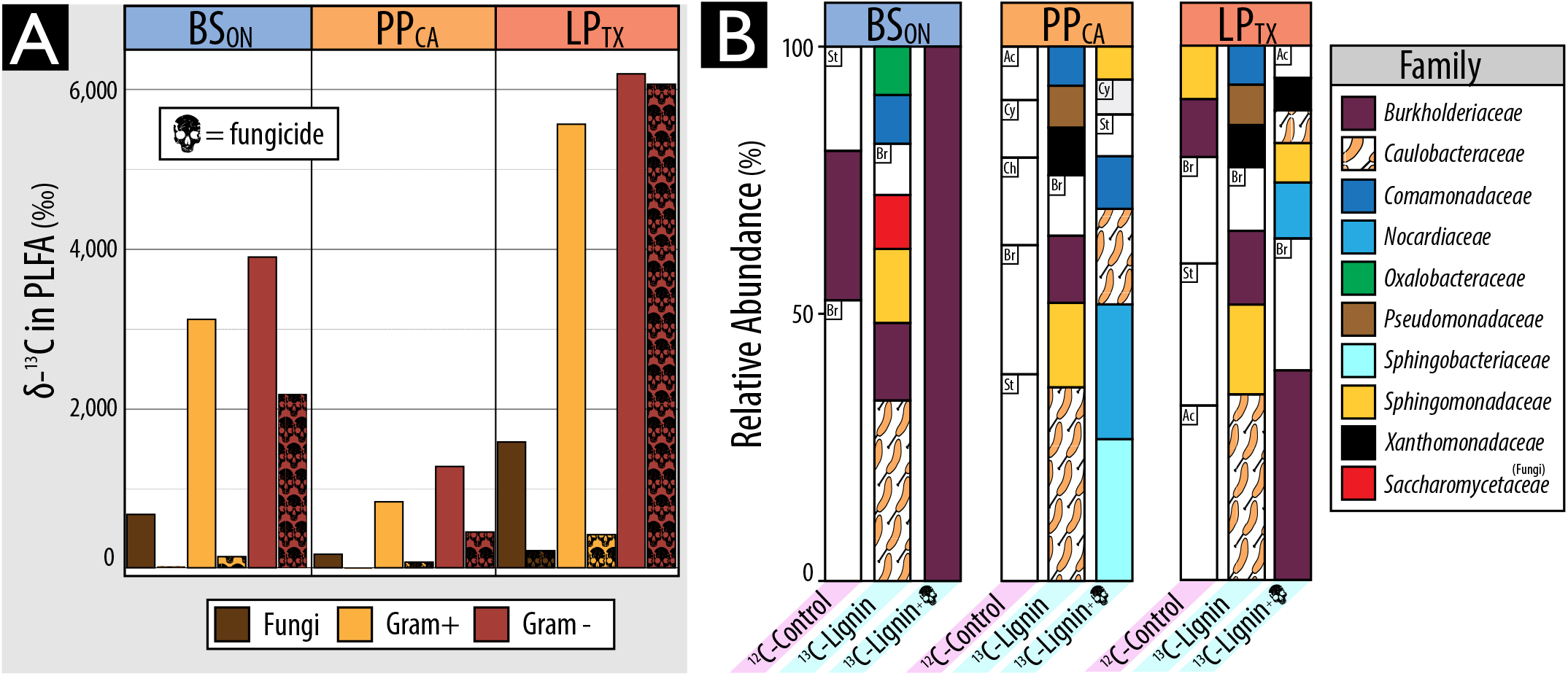
Impact of fungicide treatment on (A) the total assimilation of ^13^C from lignin in to PLFA and (B) the relative abundance of taxa in ^13^C-metagenomes. In B, BS_ON_ was averaged from two replicates. Taxa that commonly occurred in ^12^C-control libraries but were not designated as major lignolytic taxa were left uncolored (St: Streptomycetaceae, Br: Bradyrhizobiaceae; Ac: Acidobacteriaceae, Ch: Chitinophagaceae and Cy: Cytophagaceae). Taxa comprising fewer than 0.05% of total reads are not displayed.

## CAZy Gene Content in Cellulose-and Lignin-degraders

Genes encoding lignin-modifying Auxiliary Activity (AA) enzymes were approximately three-fold more abundant in ^13^C-lignin versus ^13^C-cellulose metagenomes. Yet, many AA families were also enriched in ^13^C-cellulose versus control metagenomes, which corresponded to an overall increase in reads classified as fungal (Figure 8). In contrast, the enriched AA gene families in ^13^C-lignin metagenomes (AA3, AA4, and AA6) were largely bacterial in origin. Overall, there was an 18-fold increase in fungal versus bacterial AA reads in ^13^C-cellulose metagenomes compared to a 4-fold increase in bacterial versus fungal AA reads in ^13^C-lignin metagenomes (Mann-Whitney, W = 92, *p* < 0.001). Genes encoding several oxidative enzymes implicated in the depolymerization of lignin were enriched in ^13^C-lignin metagenomes and were classified to taxa previously identified as assimilating ^13^C from lignin in amplicon libraries. More specifically, genes encoding laccases (AA1), dye-decolouring peroxidases (DyP) and alcohol oxidases were the most enriched AA gene families. These were predominantly classified to Commamonadaceae, Sphingomonadaceae and Caulobacteraceae (Figure S9a). Nine MAGs contained genes predicted to encode the entire β-ketoadipate pathway, potentially involved in catabolizing aromatic lignin depolymerization products to central metabolites, and were classified to *Burkholderia*, *Sphingomonas*, *Caulobacter* and *Sorangium.* Aryl alcohol oxidase genes were the most significantly correlated (ρ > 0.3 and p_adj_ < 0.01) with total ^13^C-assimilation from lignin into PLFA.

**Figure 8.**
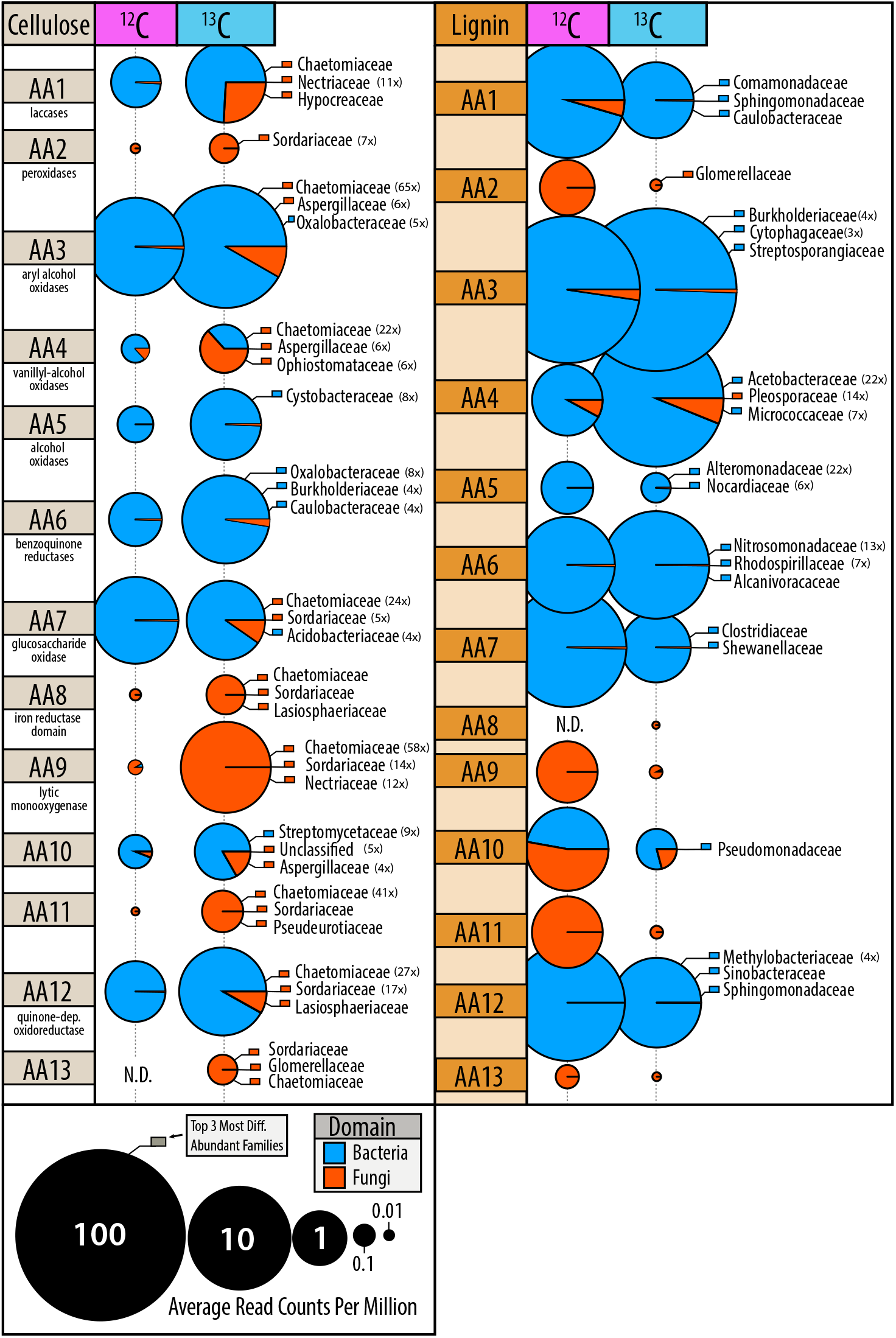
Differences in proportion of fungal and bacterial encoded auxiliary activity (lignin-modifying) genes in ^13^C-versus ^12^C-metagenomes. Bubbles are coloured according to domain of life and scaled to the total read counts per million. The top three most enriched taxa, at least 3-fold more abundant in ^13^C-libraries, are displayed and coloured according to domain of life. The fold-change of enrichment for each taxa is provided in brackets.

^13^C-cellulose metagenomes were highly enriched in glycosyl hydrolase gene families (GHs). Approximately 37% of CAZy gene clusters (groups of 3 or more GHs) from ^13^C-cellulose metagenomes contained carbohydrate-binding module (CBM) genes and 4% of these also contained an endoglucanase gene. In comparison, only 22% of clusters in ^13^C-lignin metagenomes contained a CBM gene, and none of these contained endoglucanase genes. Thirty-three percent of endoglucanase gene-containing clusters contained a CBM, and were mostly classified to *Asticcacaulis, Sorangium* and *Chthoniobacter* (Figure S9b). Actinomycetales had the greatest diversity of endoglucanase gene families in CAZy gene clusters, while fungi predominantly had endoglucanase genes belonging to GH7 and GH9 families (Figure S9c). Few endoglucanase or GH gene families were correlated with cellulolytic activity (Table S7). Four MAGs contained CAZy gene clusters encoding enzymes putatively catabolizing all three substrates (i.e. xylanases, endoglucanases and ligninases) and were classified as *Chaetomium* (fungi), *Caulobacter*, *Chthoniobacter* and *Sphingobium* (Table S8).

## Discussion

We failed to find support for our initial hypothesis that individual species would degrade multiple components of lignocellulose. Instead, we identified a diverse array of taxa assimilating ^13^C from three major lignocellulosic polymers, demonstrating the specialization of decomposer populations. While approximately one third of taxa enriched in ^13^C-DNA pools possessed members that could collectively degrade more than one substrate, and, though a handful of MAGs encoded a putative suite of catabolic enzymes, none of the individual bacterial OTUs (at a 99% similarity threshold) were definitively enriched from all three substrates. Even closely related taxa, like *Caulobacter* and *Asticcacaulis*, exhibited differences in their levels of enrichment on cellulose and lignin and in CAZy gene content. In contrast, the multi-substrate degradative capacity of fungi was evident in the increased relative abundances of genes encoding both delignifying (peroxidases) and cellulolytic enzymes in ^13^C-DNA pools. The recovery of a *Myceliophthora* MAG from ^13^C-metagenomes was consistent with its known capability for complete lignocellulose decomposition (Li *et al.*, 1999; Babot *et al.*, 2011). Therefore, our findings suggest that complementation among functional guilds may be necessary for decomposition of lignocellulose by bacteria and not necessarily fungi. One caveat is the possibility that our method was biased towards identifying single substrate utilizing bacteria. The activity of multi-substrate users may have been masked by populations that grew faster on individual substrates, particularly the relatively labile hemicellulose, as was observed in other SIP experiments (Pepe-Ranney and Campbell *et al.*, 2016). At the very least, our study demonstrates the decomposition of lignocellulosic polymers can commonly occur via a division of labour among specialized taxa.

The substantial assimilation of carbon from our model lignin substrate by bacteria supports our hypothesis that bacteria contribute significantly to degradation of native forms of lignin *in situ*. Bacterial activity was particularly evident in deeper mineral layers of forest soil, suggesting that lignin decomposition can occur throughout the soil column. Interestingly, the most novel lignin-degraders from mineral soils belong to uncultured clades of Caulobacteraceae, Acidobacteria, Solirubrobacterales, Elusimicrobia, Nevskiales, and Cystobacteraceae. The low activity of fungal lignin degraders in our microcosms was unlikely due to unmet nutritional or environmental conditions, since metabolically competent taxa like *Myceliophthora* (Li *et al.*, 1999; Babot *et al.*, 2011) assimilated carbon from cellulose under the same conditions. One possible explanation is that fungi were involved in lignin-degradation but did not metabolize degradation products to become sufficiently labeled, perhaps because of bacterial cross-feeding, evident in the inhibitory effect of fungicide on lignin incorporation by bacteria. Indeed, the results from the fungicide-treatments indicated significant interactions occurred between bacteria and fungi in lignin degradation. However, fungi were not essential for bacterial lignin degradation, given the persistent and substantial incorporation of ^13^C-lignin by bacteria in fungicide-treated soils. This study provides strong evidence for the capacity of bacteria to degrade lignin, highlighting the need for further characterization of their activity and ecology in soil.

The substantial number of taxa identified by SIP with previously reported lignocellulolytic activity was testament to the success of previous culturing-based efforts to characterize soil decomposers and a validation of our SIP approach. Approximately 72% (14/19) of hemicellulolytic taxa and 74% (31/42) of cellulolytic taxa were previously reported to degrade the corresponding substrate *in vitro.* Far fewer lignolytic taxa (~28%, 8/29) had previously reported degradative activity, likely due to less frequent and formalized testing of lignin degradation. However, there was considerable agreement in the bacterial genera we identified and those identified in other culture-independent studies of lignin-degraders in forest soil, such as *Caulobacter*, *Sphingomonas*, *Sphingobacterium*, *Nocardia*, *Telmatospirillum* and *Azospirillum* (DeAngelis *et al.*, 2011a; Woo *et al.*, 2014; Pold *et al.*, 2015). A lower proportion of lignin-degrading taxa associated with mineral layer soil (5/16) had previously reported activity compared to those associated with the organic layer (8/12), suggesting the former may be more difficult to culture or were less frequently targeted for study.

Many of the novel cellulolytic groups we identified belong to cultivation-resistant phyla, Planctomycetes, Verrucomicrobia, Chloroflexi and Armatimonadetes (formerly 0P10), which commonly predominate soil communities (Youssef *et al.*, 2009; Bergmann *et al.*, 2011). Each of these phyla possesses at least one isolate capable of degrading cellulose (Sangwan *et al.*, 2004; Lladó *et al.*, 2015; Dedysh *et al.*, 2013; Lee *et al.*, 2014) and have been designated cellulolytic in other SIP-cellulose experiments Eichorst and Kuske, 2012; Verastegui *et al.*, 2014; Pepe-Ranney and Campbell *et al.*, 2016; Wilhelm^a^ *et al.*, 2017). Notably, all contigs that contained clusters of ten or more CAZymes (27 contigs), from ^13^C-cellulose metagenomes, were classified to taxa from the aforementioned groups and contained both CBMs (26/27) and endoglucanases (20/27). These findings supported our assertion that the richness of described lignocellulolytic taxa is underestimated in soils, which limits our understanding of the ecology of decomposition and may afford new types of biocatalysts for processing lignocellulosic biomass.

The widespread capacity among members of Caulobacteraceae to degrade the three components of lignocellulose was unexpected, yet consistent with reports of their enrichment on decaying wood (Folman *et al.*, 2008; Valaskova *et al.*, 2009) and during early stages of litter decomposition (Smenderovac, 2014). *Caulobacter* were first isolated from cellulose-amended lake-water (Henrici and Johnson, 1935), but are primarily known as oligotrophic, aquatic organisms (Poindexter *et al.*, 1981). Their role in degradation of plant carbohydrates was first postulated based on the analysis of the *C. crescentus* genome (Nierman *et al.*, 2001) and subsequently demonstrated by growth on cellulose, xylose and vanillate (Hottes *et al.*, 2004; Thanbichler *et al.*, 2007; Song *et al.*, 2013; Presley *et al.*, 2014). Although we provide the first evidence for the role of Caulobacteraceae in degrading all three polymers of lignocellulose, several surveys of forest soils report enrichment of *Caulobacter* or *Asticcacaulis* in samples amended with cellulose (Verastegui *et al.*, 2014; Wang *et al.*, 2015; Pepe-Ranney and Campbell *et al.*, 2016) or lignin (DeAngelis *et al.*, 2011; Pold *et al.*, 2015). The role of Caulobacteraceae in decomposition in forest soils may prove significant, given their relatively high abundance (0.5 – 2.5% of total libraries) and their capacity to adhere to insoluble polymers, like lignocellulose.

The taxonomy and catabolic capacity of lignocellulose degraders were moderately associated with variation in the total lignocellulolytic activity. The influence of compositional differences between soil layers was, in certain cases, overshadowed by the strong negative correlation between organic matter and ^13^C enrichment. Yet, in regression modeling, CAZy gene content and community composition had equivalent or greater explanatory power than other predictors. The abundances of several prominent lignocellulolytic taxa were significantly correlated with higher ^13^C assimilation, demonstrating that certain taxa play more significant roles, at least under conditions we tested. Strongly oxidative bacterial enzymes commonly studied in relation to lignin-degradation, such as DyP-type peroxidases and laccases (Ausec *et al.*, 2012; Colpa *et al.*, 2014; Singh and Eltis, 2015), were less predictive of lignolytic activity than aryl alcohol oxidases, suggesting a greater role of the latter class of enzymes in bacterial lignin-degradation. Overall, fewer endoglucanase genes were predictive of ^13^C-assimilation compared to AA gene families. This difference suggests either the existence of a greater diversity of endoglucanases, minimizing the explanatory power of any single gene, or a narrower activity range of AA families with a correspondingly higher relevance to lignolytic activity. This result agrees with the finding that the composition of decomposer communities had increased explanatory power for the rate of litter decomposition during later stages when greater proportions of lignin remained (Cleveland *et al.*, 2014).

The comprehensive study of all three major lignocellulosic polymers enabled us to examine the co-occurrence of lignocellulolytic traits, addressing several knowledge gaps in this area. The results indicated unexpected specialization among bacterial populations for degradation of individual lignocellulosic polymers and revealed several novel lignocellulolytic taxa, highlighting current limits to our knowledge of decomposition. This research supports the view that bacterial decomposition of oligomeric lignin is a ubiquitous soil process, with the potential to occur in deeper soil layers following early stages of litter decomposition. As hypothesized, variation in community composition was found to constrain lignocellulolytic activity across various forest types and soil layers in North America. The relationship between these communities and process rates should receive continuing study to refine our understanding of soil carbon stabilization and terrestrial carbon cycling models. Furthermore, the large number of degradative gene clusters from uncultivated lignocellulolytic taxa represent a trove of potentially novel enzymes for biotechnological applications.

## Acknowledgements

This study was supported by a Large Scale Applied Research Project (162MIC) funded by Genome Canada and Genome BC. R. Wilhelm was supported by a NSERC graduate scholarship and UBC Four Year Fellowship.

## Conflict of Interest

The authors declare no conflict of interest.

---

**Figure S1.**
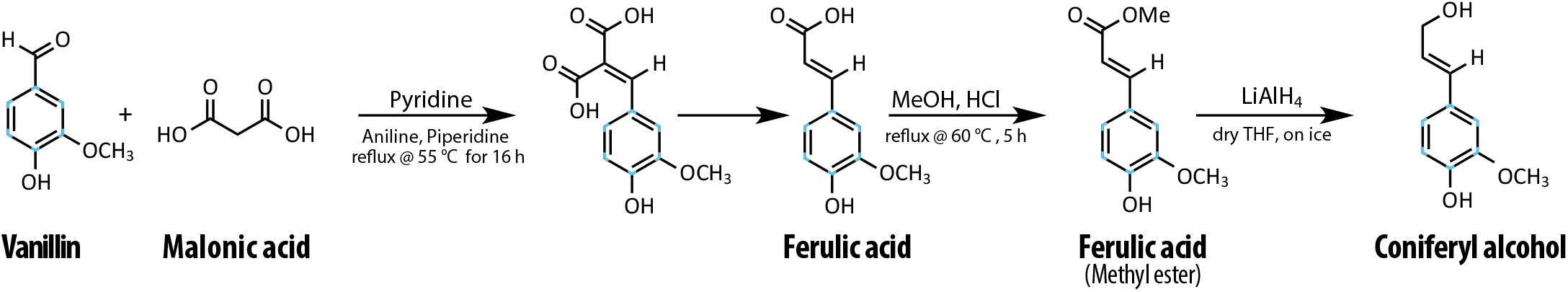
Reaction scheme for the synthesis of ^13^C ring-labeled coniferyl alcohol which was subsequently polymerized into lignin with an average of fourteen coniferyl alcohols. Carbon-13 isotopes are denoted in light blue. Consult the Supplementary Methods for a complete description of synthesis reactions.

**Figure S2.**
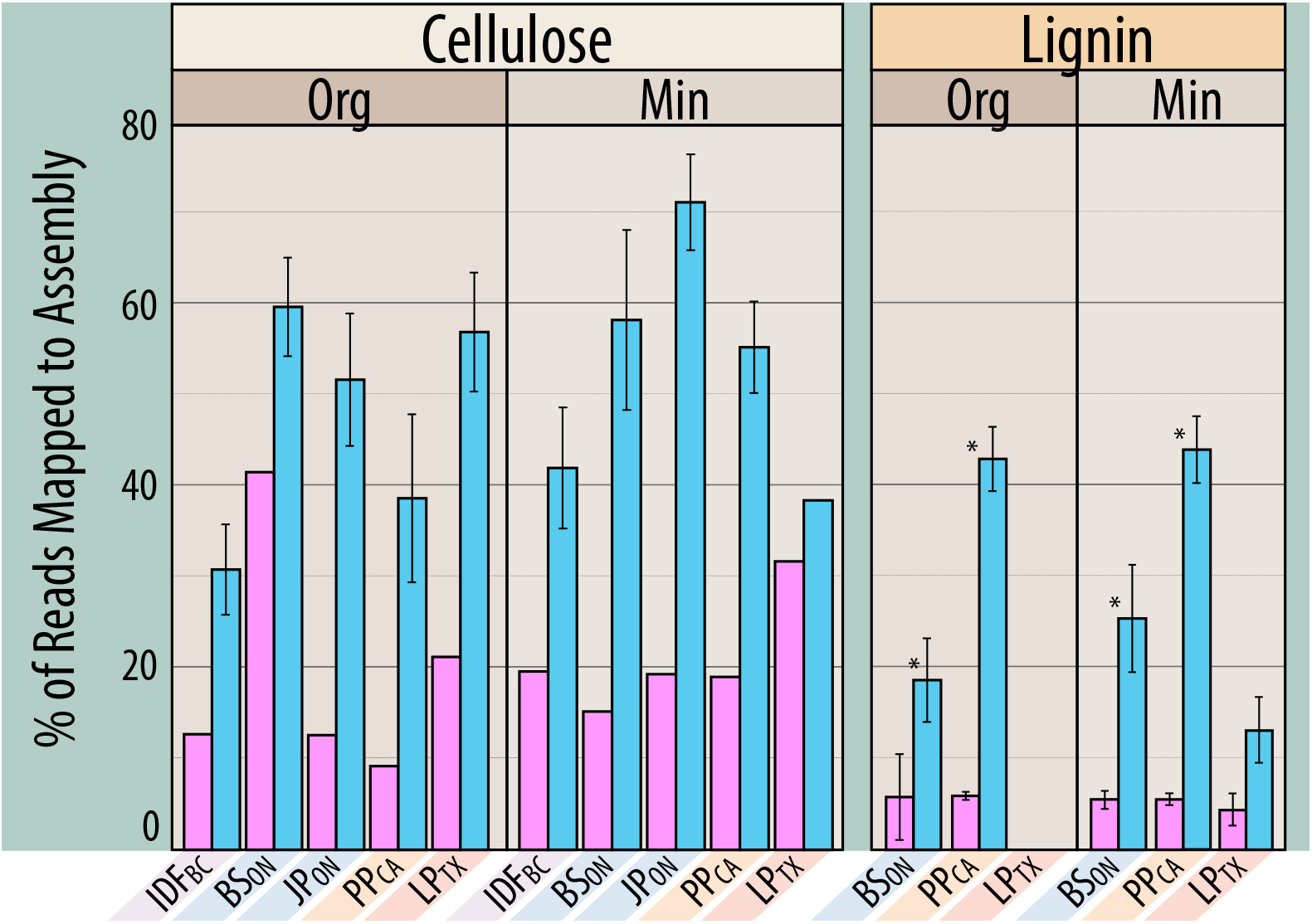
Evidence for the improved assembly of metagenomes derived from ^13^C-en-riched DNA in soilmicrocosms amended with ^13^C-labeled (blue) or unlabeled (pink) cellulose or lignin. Statistically supported differences between paired labeled and unlabeled treatments (t-test; p < 0.01) are designated with an asterix (*). Statistical testing could not be performed for cellulose with single ^12^C-libraries. When composited, ^12^C-libraries cellulose libraries had significantly lower assembly than ^13^C-libraries in organic (Wilcoxon; p < 0.003) and mineral layer soils (Wilcoxon; p < 0.001).

**Figure S3.**
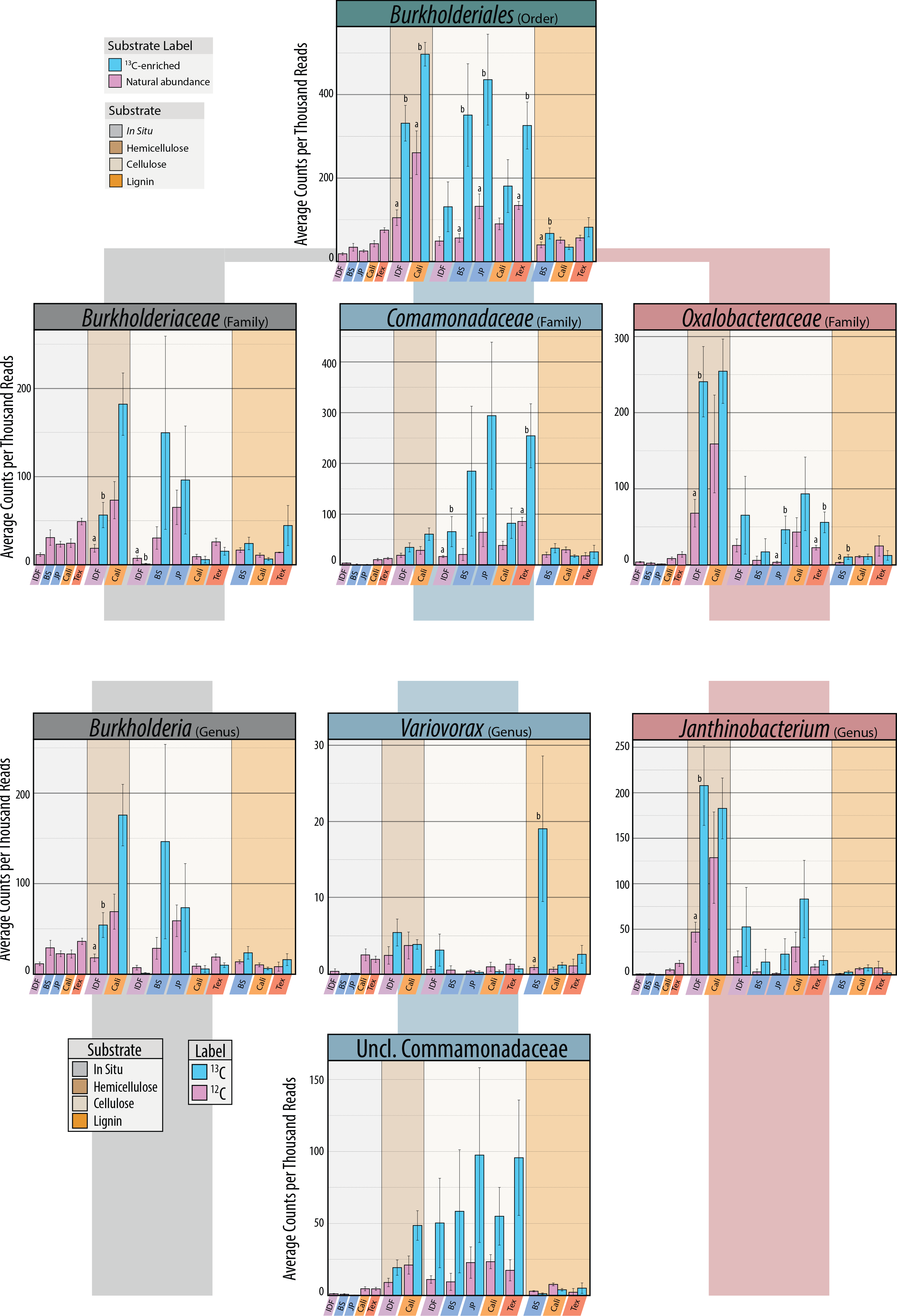
The relative abundance of prominent lignocellulolytic members of Burkholderiales based on differential abundance between ^12^C-and ^13^C-libraries. Plots are organized in a hierarchical structure displaying family and genus for all active taxa within each group. Error bars correspond to one standard error of the mean. Significant (Tukey HSD; p_adj_ < 0.05) pairwise differences are grouped by lettering.

**Figure S4.**
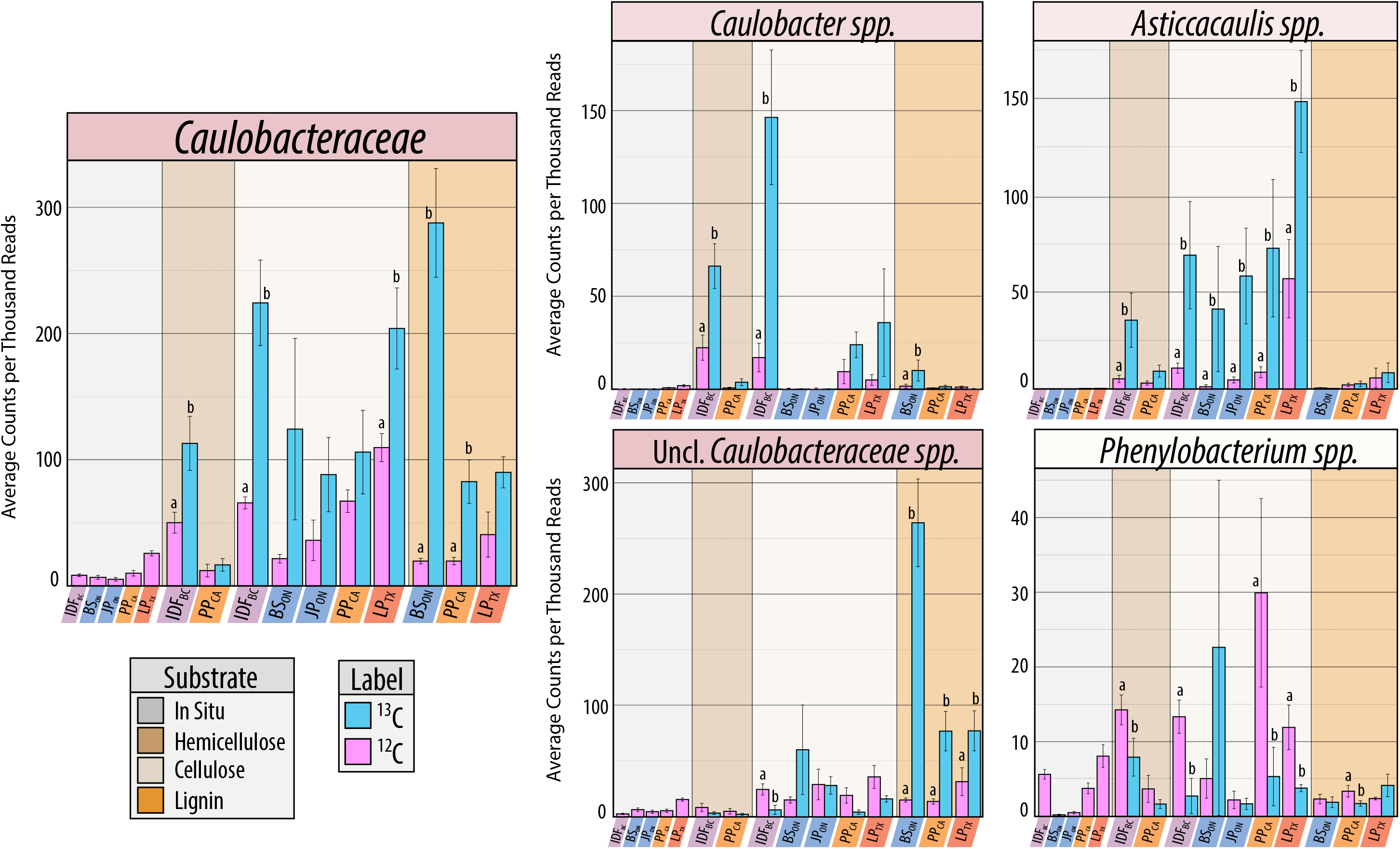
The relative abundance of all OTUs classified to the genera of Caulobacteraceae. Differences in hemicellulolytic, cellulolytic and lignolytic can be observed among genera and unclassified Caulobacteraceae sequences based on differential abundance between ^12^C-and ^13^C-libraries. Error bars correspond to one standard error of the mean. Significant (Tukey HSD; p_adj_ < 0.05) pairwise differences are grouped by lettering.

**Figure S5.**
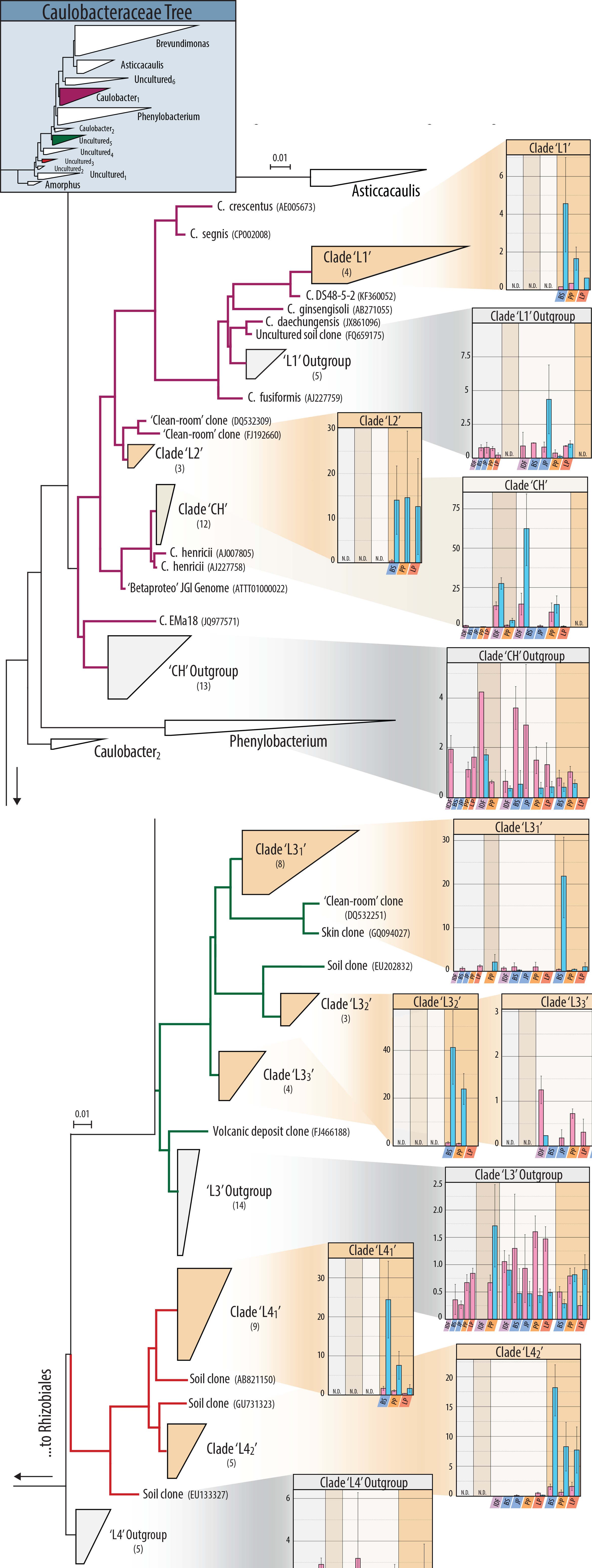
Maximum parsimony tree showing phylogenetic distribution of lignocellulolytic OTUs within the familyCaulo-bacteraceae. Clades are named based on SILVA tree and custom names were assigned to putatively functional clades. Branches were coloured to indicate membership to broader clades inset at the top left. The closest cultured represtatives were included where possible. Clades from this study are comprised of multi-ple OTUs with the total number provided in parentheses.

**Figure S6.**
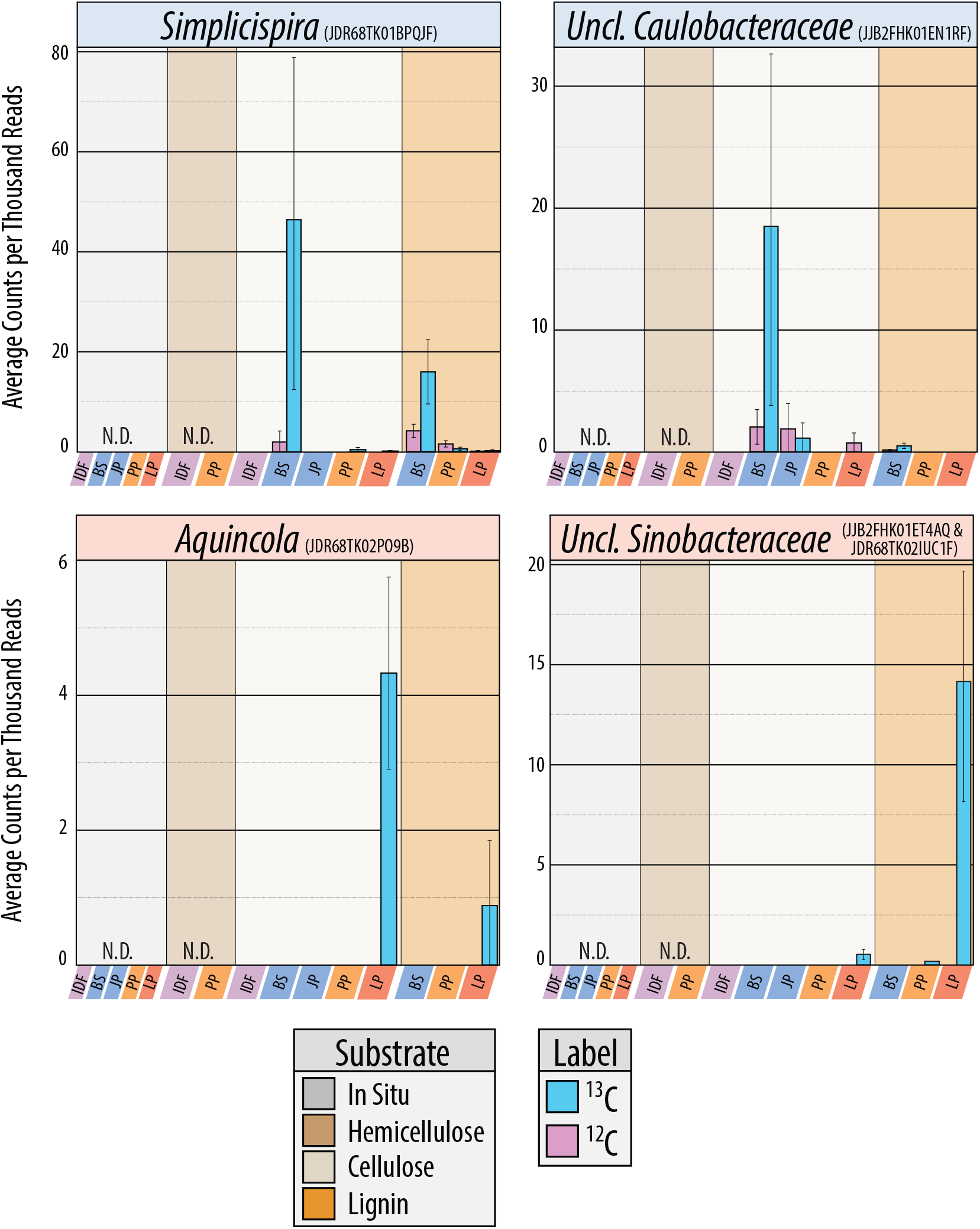
Abundances of OTUs enriched in both lignin and cellulose ^13^C-pyrotag libraries within the same region. Plots are titled with the lowest supported taxonomic classification and include representative read names. Where two OTUs were combined, both independently exhibited the trend in multiple substrate use. The unclassified Caulobacteraceae OTU belongs to the clade ‘LH3_1’ presented in Supplementary Figure 3b.

**Figure S7.**
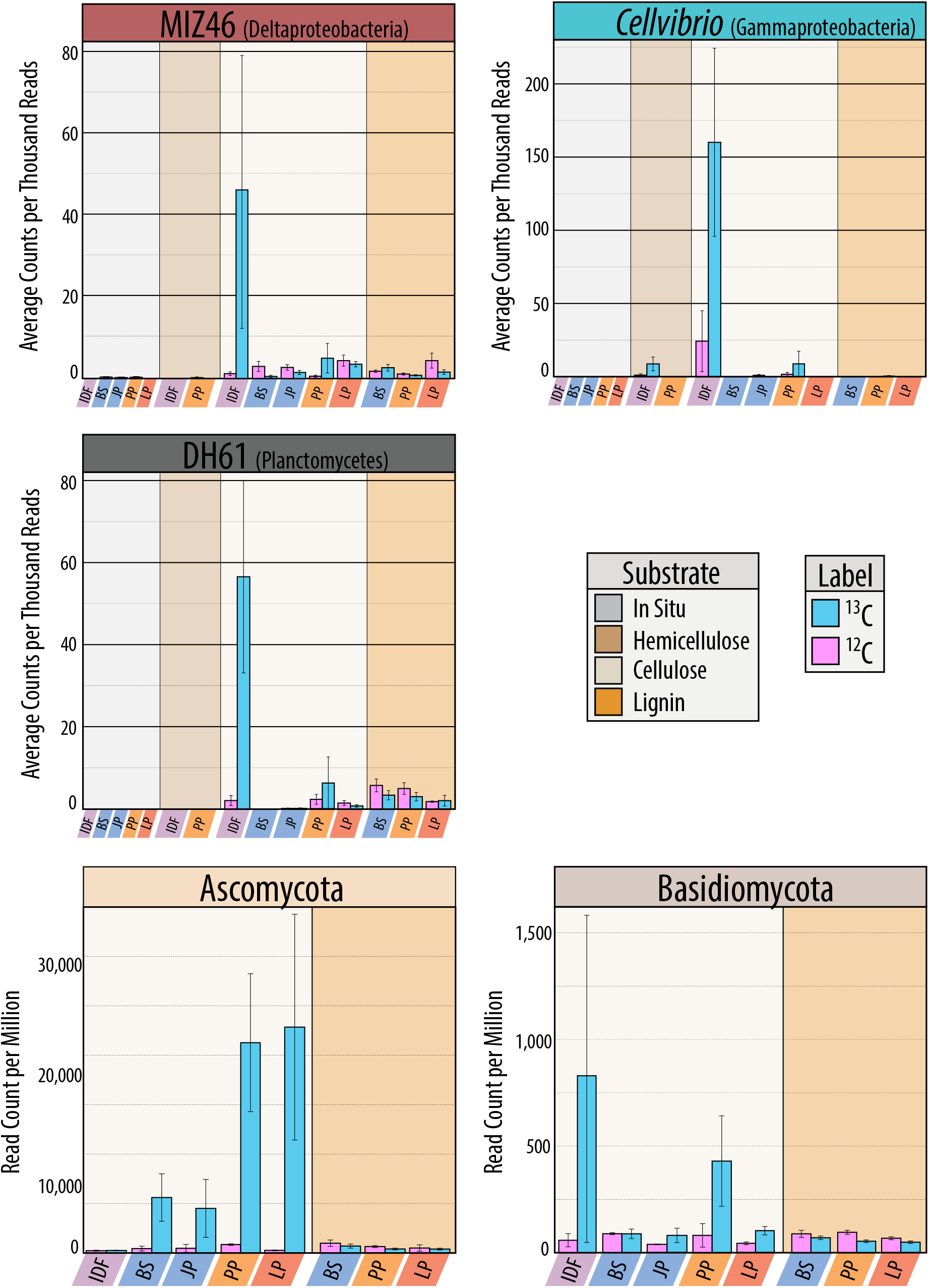
Relative abundances of cellulolytic taxa unique to IDF_BC_.The relative abundances of bacteria are based on 16S rRNA gene pyrotag libraries and fung on LCA classification of unassembled shotgun metagenomes. Error bars correspond to one standard error of the mean.

**Figure S8.**
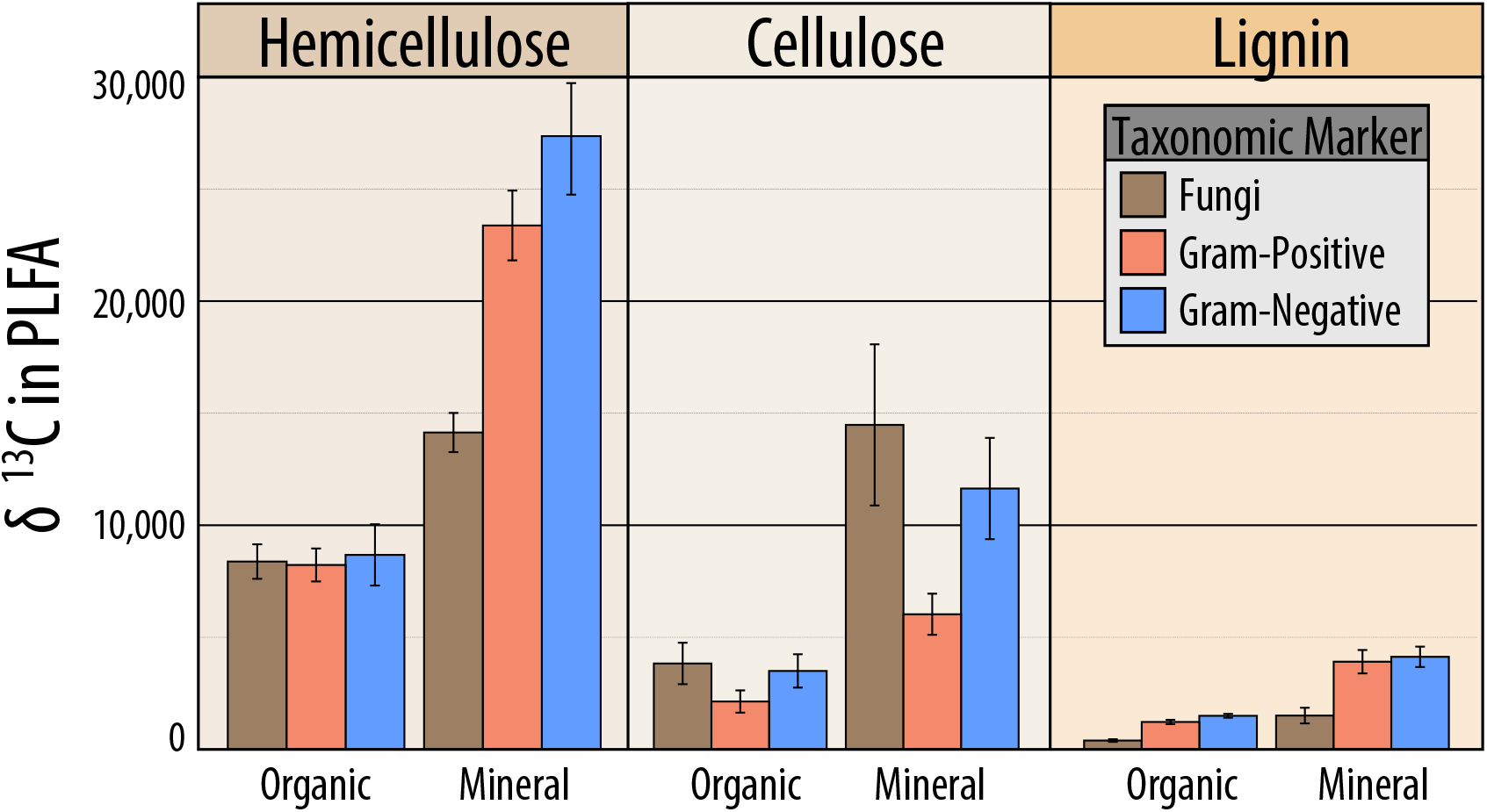
The average delta-^13^C enrichment of PLFA markers for fungi, gram-positive bacteria and gram-negative bacteria for all three substrate amendments.

**Figure S9.**
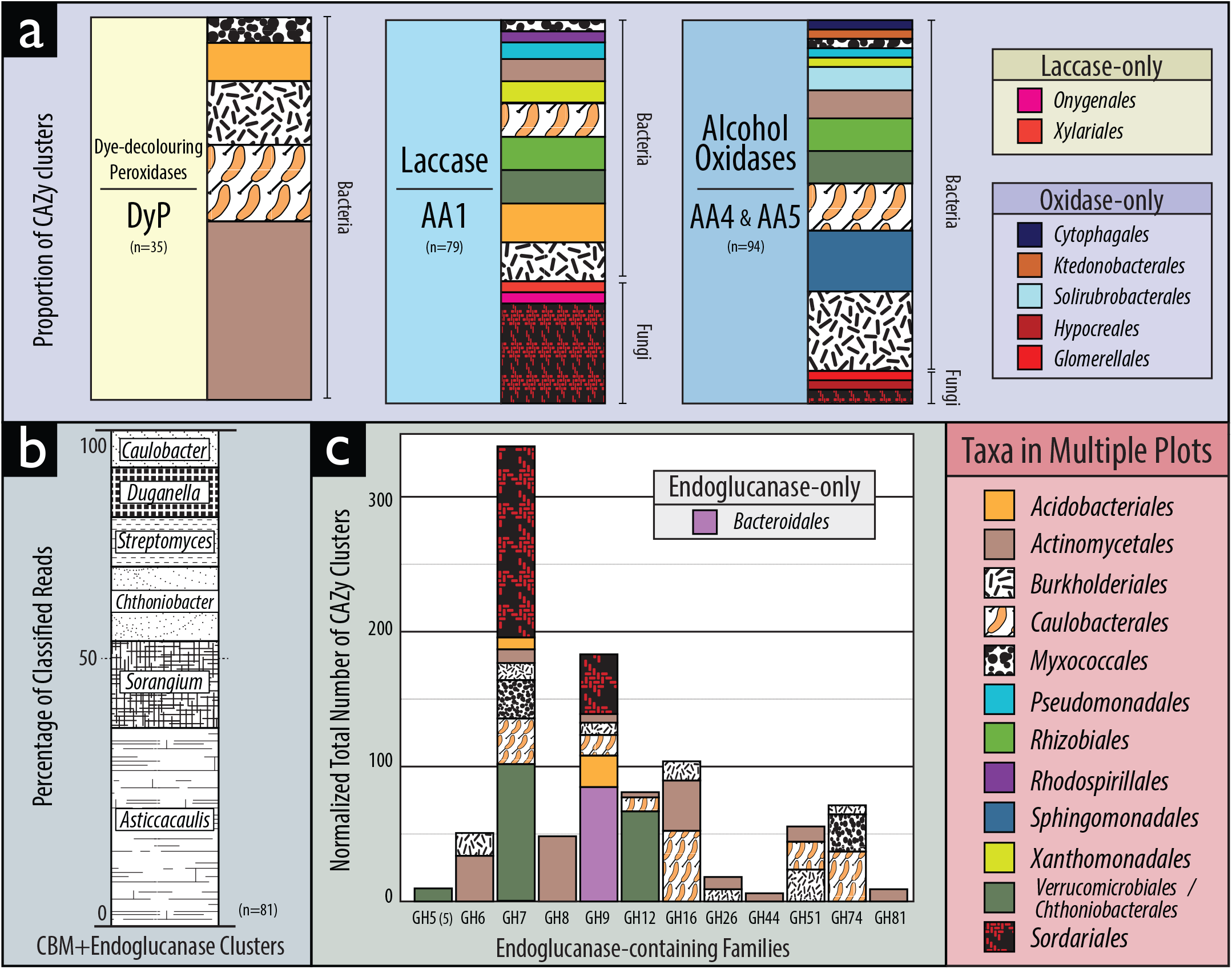
Taxonomic classification of catabolic genes in CAZy clusters in ^13^C-metagenome assemblies, including (A) genes encoding lignin-modifying enzymes, (B) clusters containing a CBM and an endoglucanase, and (C) all endoglucanase-containing gene families. All taxa represented by a single gene were not displayed. Taxonomic classifications were assigned based on LCA and functional annotation based on top blast hit (e-value < 10^−5^) to CAZy database or positive match to custom hidden Markov models.

